# Characterization of the oral microbiome of medically controlled type-2 diabetes patients

**DOI:** 10.1101/2020.04.07.031070

**Authors:** Ana Almeida-Santos, Daniela Martins-Mendes, Magdalena Gayà-Vidal, Lucía Pérez-Pardal, Albano Beja-Pereira

## Abstract

Type 2 diabetes mellitus (T2DM) is a chronic metabolic disease that is becoming a significant global health care problem. Several studies have shown that people with diabetes are more susceptible to oral problems, such as periodontitis and, although the causes are still inconclusive, the oral microbiota seems to be an important factor in this interaction. This study aimed to characterize the oral microbiome of a sample representing T2DM patients from Portugal and exploit potential associations between some microorganisms and variables like teeth brushing, smoking habits, and nutrient intake. By sequencing the hypervariable regions V3-V4 of the 16S rRNA gene in 50 individuals belonging to a group of diabetes patients and a control group, we found a total of 233 taxa, from which only 65% were shared between both groups. No differences were found in terms of alpha and beta diversity between groups or habits categories. Also, there were no significant differences in the oral microbiome profiles of control and diabetes patients. Only the class *Synergistia* and the genus *TG5*, which are related to periodontitis, were statistically more frequent in the control group. This finding can be justified by the fact that these diabetic patients usually have their oral health under close medical control than an average healthy person, which in this study was represented by the control group.

**IMPORTANCE:** Diabetes has become a significant global health care issue as its incidence continues to increase exponentially, with type 2 diabetes being responsible for more than 90% of these cases. Portugal is one of the countries with a higher prevalence of diabetes in Europe. It has been reported that diabetic people have an increased risk of developing several health problems such as oral infections mostly caused by opportunistic pathogens. Some studies have pointed out a relationship between diabetes and oral microbiome. Therefore, the characterization of the microbial ecosystem of the mouth in reference groups is crucial to provide information to tackle oral health pathogen-borne conditions. In this study, we provide the first characterization of the oral microbiome of type 2 diabetes mellitus patients from Portugal, and therefore, contributing new data and knowledge to elucidate the relationship between diabetes and the oral microbiome.

## Introduction

Type 2 diabetes mellitus (T2DM) is a metabolic disease characterized by chronic hyperglycemia caused by defects in insulin secretion, which can contribute to the development of resistance to its action (1, 2). T2DM is becoming more common, also in children and adolescents, and therefore, represents a significant global health care problem (3).

As more studies are focused on the impact of T2DM on the medium-term patients’ health, a growing number of studies have been reporting a close association between diabetes and susceptibility for some oral illnesses, such as periodontitis (2), derived from the deregulation of the oral microbiota equilibrium that increments the establishment of pathogenic organisms, which leads to the presence of oral pathogenic bacteria, causing the deregulation of the oral microbiota equilibrium, and vice-versa.

The oral microbiota is among the most diverse and dynamic ecosystems of the human body, in which it has been identified more than 700 species of bacteria (4). Bacterial phyla *Firmicutes, Actinobacteria, Fusobacteria, Proteobacteria*, and *Bacteroidetes* dominate the oral microbiota, accounting for 80-95 % of the total species (5). These microorganisms normally harmoniously co-exist with their host due to coevolution, however, behavioral factors such as poor oral hygiene and diet, debilitated immune systems, genetics, medication and, certain diseases can lead to a dysbiotic oral ecosystem (6). This microbial imbalance is normally associated with an overgrowth of pathogenic microorganisms, which can lead to more susceptibility to oral illness (7, 8).

The oral microbiota plays an important role in the relationship between periodontitis and diabetes (9) since it influences glycemic control (10). Certain bacteria such as *Porphyromonas gingivalis*, one of the main strains of periodontal disease, releases a lipopolysaccharide (LPS), which triggers numerous pro-inflammatory cytokines that are responsible for periodontal tissue destruction as well as increases insulin resistance (1, 11, 12). However, other taxa could be related to T2DM. For example, J. Long et al. (4) compared the oral microbiome profiles from African Americans subjects with T2DM with non-diabetic obese individuals and non-diabetics with normal-weight and found a higher abundance of taxa in the phylum *Actinobacteria*, which were associated with a lower diabetes risk; whereas *Actinomycetaceae, Bifidobacteriaceae, Coriobacteriaceae, Corynebacteriaceae, and Micrococcaceae* families were less abundant among diabetic subjects compared to normal-weight controls. Additionally, another study found a decrease in the biological and phylogenetic oral microbiome diversity in diabetics in comparison to non-diabetics from South Arabia, evidence that was related to an increase in the pathogenic content in the diabetic’s oral microbiome (13).

Presently, only two studies on the relationship between type 2 Diabetes Mellitus and the oral microbiome were made on European populations (14, 15). However, one of them was focused on the subgingival microbiome (14), and the other one on obese T2DM (15), and both of them considered a limited sample size (n < 20). Since the oral microbiome is an important factor for the maintenance of human health, more studies are needed to better understand the relationship between diabetes and oral microbiome.

In this study, we characterized the oral microbiome of a group of medically controlled type 2 diabetes patients from Portugal and compared them to a control group, by using amplicon sequencing of the hypervariable V3-V4 regions of the 16s rRNA from saliva samples of 50 individuals.

## Results

### Sequencing data and taxonomic assignment

A total of 12,754,645 raw reads were obtained with a mean of 255,092.9 reads per sample (ranging from 147 to 1,085,170 reads per sample). After quality filtering and mitochondria and chloroplast removal, 752,526 reads remained for further analyses.

Considering the 46 samples included in the analysis, the high-quality reads were assigned to 10,746 ASVs with a total absolute frequency of 605,211. The mean ASVs frequency per sample was 12,351.24 (range: 1,973–35,935). The ASVs were assigned to 233 taxa (Table S3). The taxa identification was possible for 71% of the ASVs at the genus level, and 21% at the species level. Control and diabetes groups shared 153 out of the 233 taxa. The control group exhibited 50 taxa that were not present in the diabetes group and the diabetes group presented 31 taxa that were not in the control group.

### Microbiome characterization

We detected a total of 13 phyla, 21 classes, 37 orders, 60 families, 86 genera and 51 species. At the phylum level, the oral cavity of all the 46 samples was dominated by Firmicutes (45%), Bacteroidetes (22%), Proteobacteria (16%), Actinobacteria (9%) and Fusobacteria (6%), constituting 98% of the total oral microbiome.

At the class level, each individual exhibited an average of 14 taxa. The ten most frequent classes accounted for approximately 97% of the total abundance in both the control and diabetes groups (Fig. S1). *Bacilli* and *Bacteroidia* are the dominant classes in both groups, accounting for 50%. *Gammaproteobacteria* had a higher abundance in the control group (9.9%) than in the diabetes group (5.4%) (Fig. S1), although this difference is not significant. Five classes were significantly different between the two groups, *Betaproteobacteria* was higher in the Diabetics group (p=0.033), and *Deltaproteobacteria* (p=0.013), Spirochaetes (p=0.035), Mollicutes (p=0.043) and *Synergistia* (p<0.001) were higher in the control group, nevertheless, after Bonferroni correction, only *Synergistia* class remained significantly different between groups (Table S2).

When focusing on the taxa abundance at the genus level, each individual displayed an average of 32 genera. The frequency distribution of the 10 most abundant taxa, up to genera is shown in Fig. 1. *Streptococcus* (29%) is the most abundant genus, present in all subjects, followed by *Prevotella* (14%) and *Neisseria* (5%).

**Figure 1.**
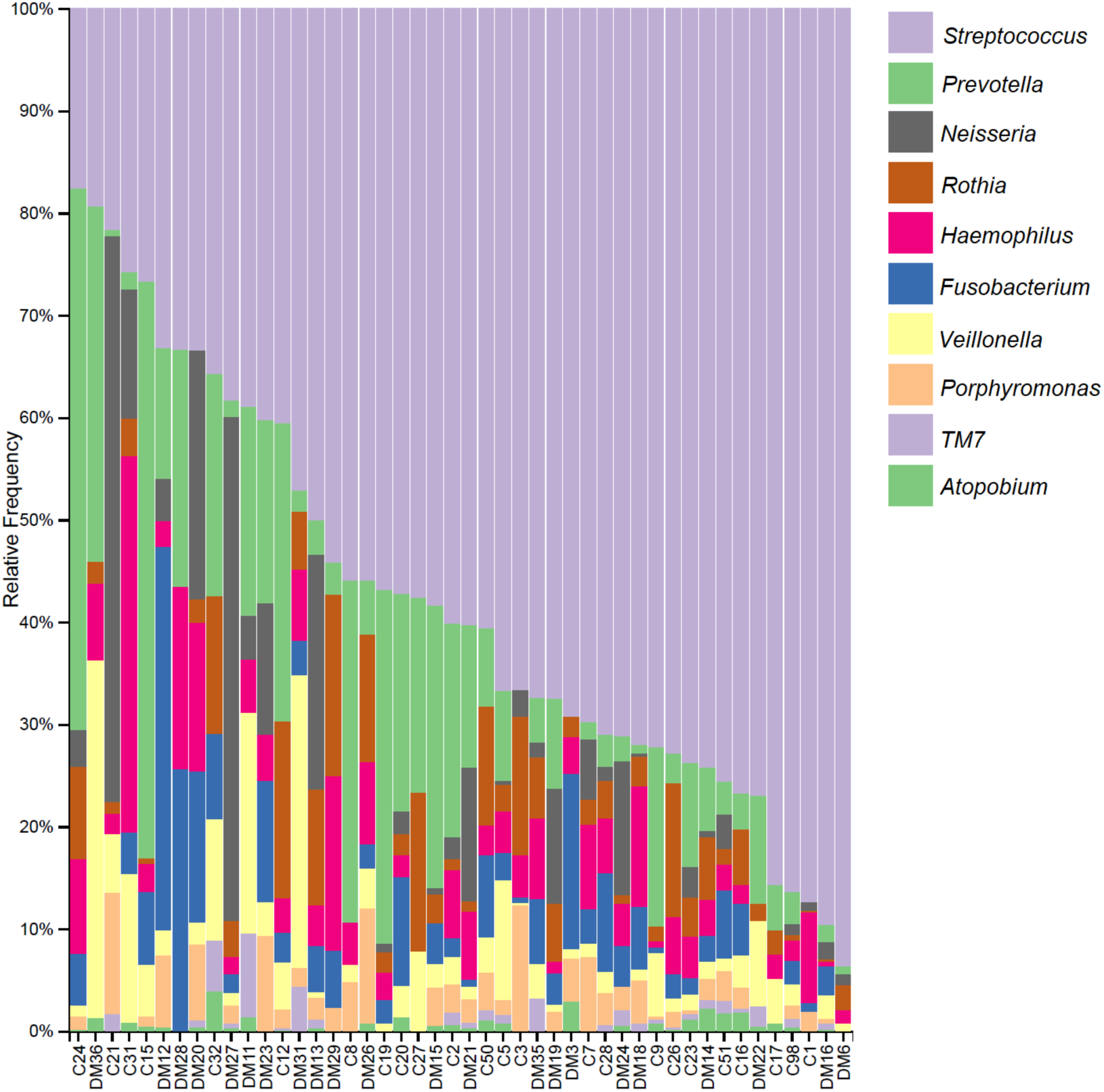
Frequency of the ten most abundant taxa up to the genus level per subject. Sample names starting with C are from the control group and those starting with DM are from the diabetics group.

The 10 most frequent genera accounted for approximately 77% in both the control and diabetes groups (Fig. 2A). It is worth noting that for some of the taxa we were able to identify them only up to the family level due to lack of resolution.

**Figure 2.**
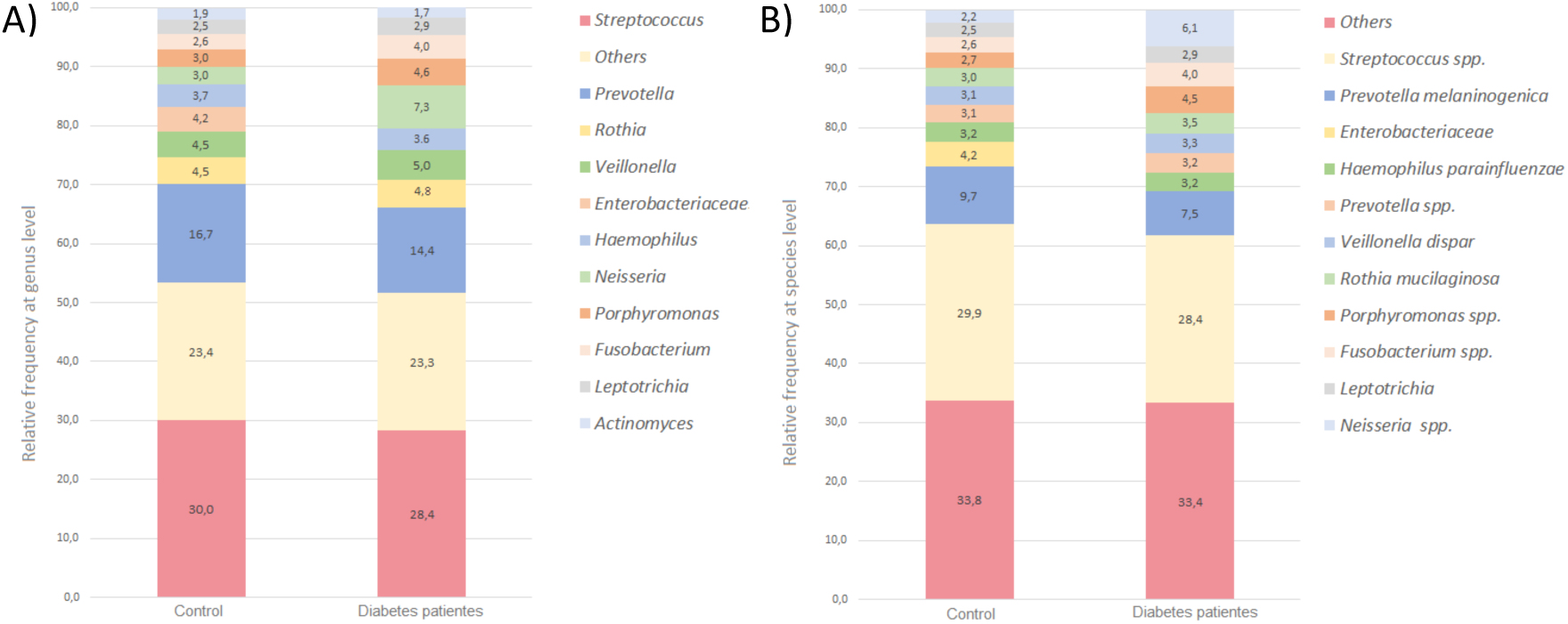
The relative abundance of the ten most-abundant taxa found at the genus level (A) and at the species level (B) in both control and diabetes groups. The remaining taxa are included in the category “Others”.

A total of 14 taxa were statistically different between the control and diabetes groups (Table S2). Nonetheless, after the Bonferroni correction, only the *TG5* genus remained significant. *TG5* genus belongs to the *Synergistia* class, the only class that was statistically significant.

At the species level, an average number of 16 different taxa per individual was observed. Note that due to the lack of resolution most taxa were not identified up to species level. The 10 most-frequent taxa (only four of them were assigned to their species level), accounted for 67% in both the control and diabetes groups (Fig. 2B). *Streptococcus spp.* and *Prevotella melaninogenica* were the two most frequent species in both groups.

A total of 20 taxa, at the species level, were significantly different between the two studied groups (Table S2), but after Bonferroni correction, only *TG5 spp*. remained statistically significant, being more abundant in the control group. The ANCOM analysis of ASV differential abundance also revealed a significantly higher abundance of the *TG5* genus and *TG5 spp.* in the control group (p <0.05).

### Diversity measures

#### Alpha diversity

The estimated Shannon index calculated with the rarefied data value was 7.0.3 ± 0.75 for the control group and 6.97 ± 0.66 for the diabetes group, whereas the ASVs abundance was 306 ± 111 and 271 ± 86 for the control and the diabetes group, respectively. We did not find significant differences in the distribution of the Shannon index nor the ASVs abundance of both groups.

#### Beta diversity

Bray-Curtis dissimilarity values were calculated to measure the differences between individuals, in terms of taxonomic structure. Figure 3 shows the distribution of the pairwise beta diversity values between individuals within each group studied and between them, and we can observe similar distributions. This was confirmed with the PERMANOVA analysis (pseudo-F=1.092; p=0.231) indicating that there is no differentiation in the microbiome composition of the diabetes group compared to the control one.

**Figure 3.**
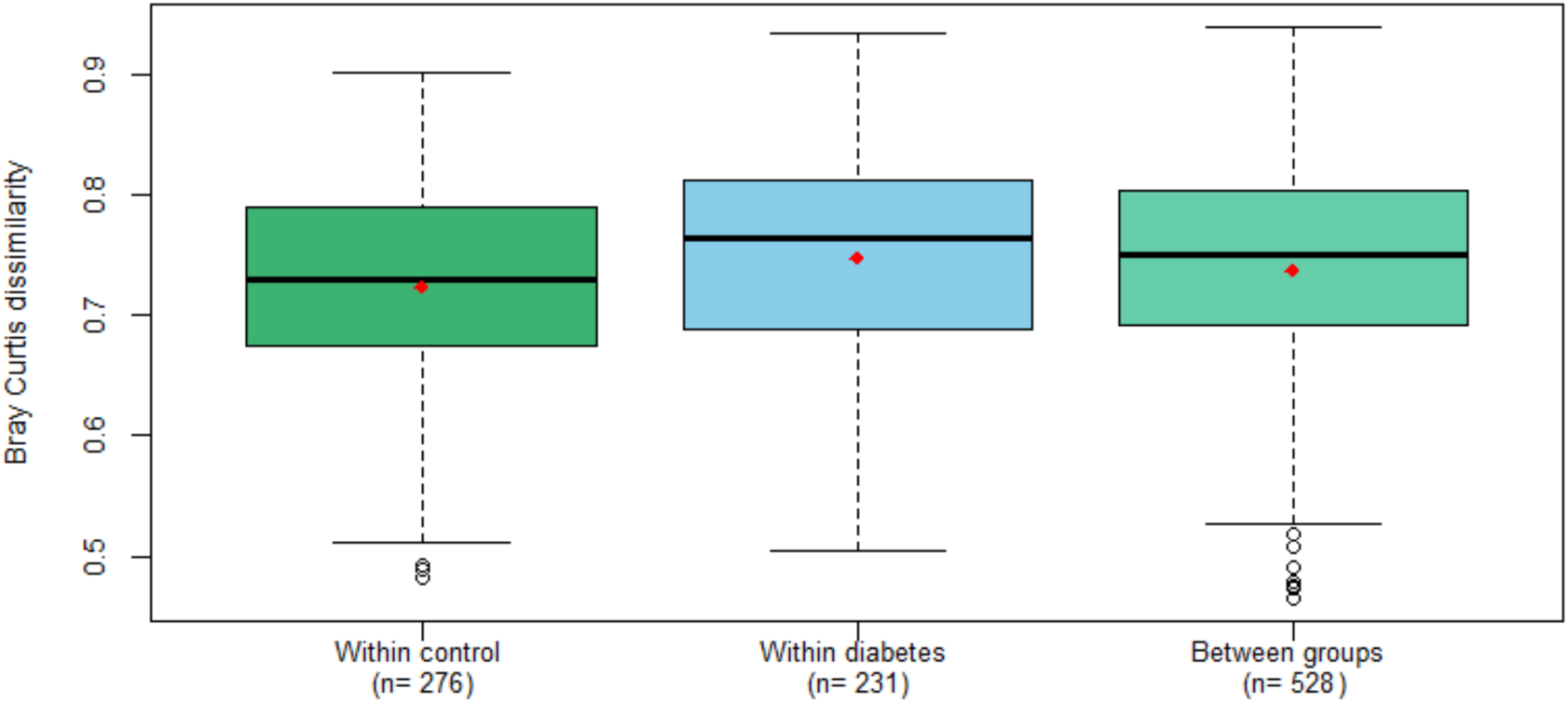
Distribution of pairwise Bray-Curtis dissimilarity values between individuals within diabetics and controls and between them. The red color dots represent the mean of each distribution.

To better visualize the differences between the microbiome profiles of the individuals analyzed, we performed a Principal-coordinates analysis (PCoA) from the Bray Curtis dissimilarity matrix that did not reveal a clear clustering pattern, being all control and diabetics individuals scattered throughout the plot (Fig. 4). The hierarchical clustering analysis formed two major clusters (Fig. S2), which seems to be due to *Streptococcus* frequency, in which the first cluster aggregates the individuals with a high frequency of *Streptococcus* and the other cluster aggregates those with lower frequencies. Both clusters are composed of individuals from the control and diabetes groups.

**Figure 4.**
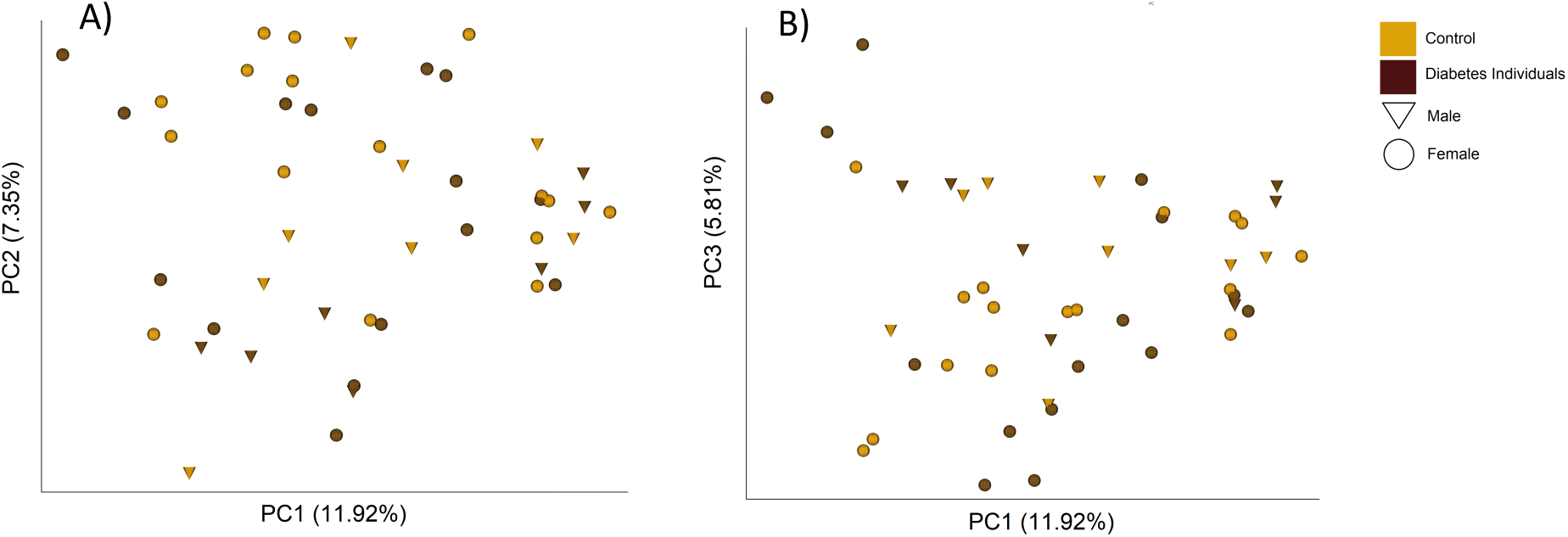
PCoA plots showing the A) first and second, and B) first and third principal components and the percentage of the total variance that they explain based on the Bray Curtis dissimilarity matrix. Each point represents one individual, with color and symbol indicating the group of study and sex.

#### Bacteria associated with periodontal disease

Due to the relationship between Diabetes and periodontal disease, we compared the relative abundance of species that have been related to periodontal disease (*Prevotella intermedia, Campylobacter rectus, Porphyromonas endodontalis*, and *Treponema socranskii* and *TG5 spp*. (16-19)), and they were almost vestigial in both groups. *TG5 spp.* was the only species statistically different between groups after the Bonferroni correction, being more abundant in the control group (Table S2).

#### Teeth brushing and smoking habits

We evaluated how the teeth brushing and smoking habits affected the oral microbiome of the individuals studied. Regarding the alpha diversity (Fig. S3), our results showed that individuals who wash their teeth once to three times per week have a higher Shannon index and ASVs abundance followed by those who wash once per day and more than once per day. Those that never wash their teeth showed the lowest index values. Concerning smoking habits, heavy smokers showed a higher Shannon index, as well as ASVs, whereas the moderate smokers presented the lowest values of both diversities’ measures. However, none of the comparisons between the above-mentioned categories were statistically significant.

The PCoA plot colored by the teeth brushing and smoking habits’ categories (not shown) did not reveal a clear clustering pattern, being all control and diabetics individuals dispersed throughout the plot. Likewise, none of the individual aggregation in the hierarchical clustering was attributable to a particular habit.

#### Diet

We collected information about nutrient intake to be able to control possible differences in the oral microbiome of diabetes and controls that could be related to diet. However, we did not find significant differences regarding nutrients consumption and energy intake between both groups after Bonferroni correction (Table 1).

**Table 1.**
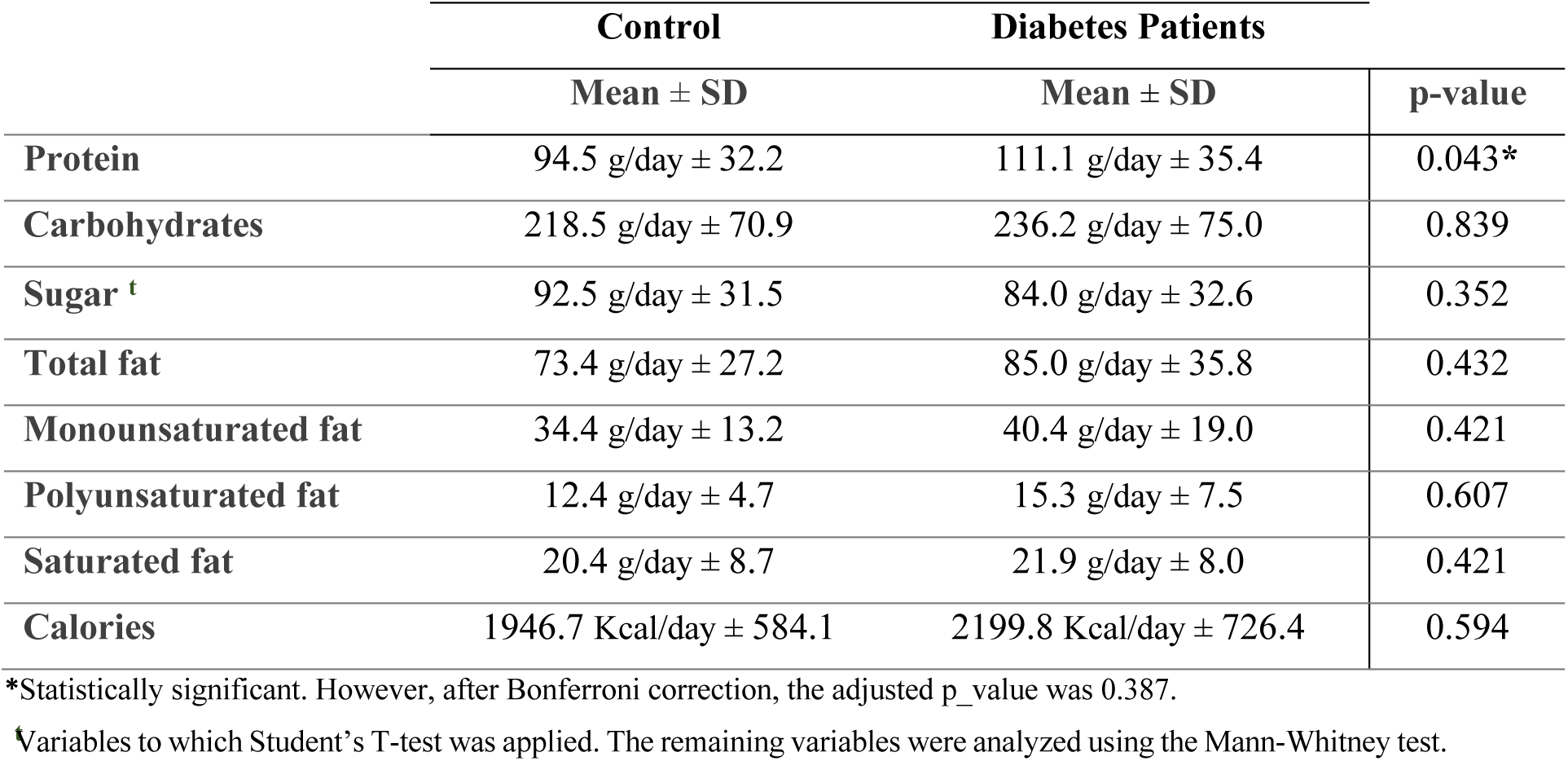
Descriptive statistics of nutrients consumption and energy intake of both control and diabetes groups and respective p-values of Mann-Whitney test/Student’s T-test between the same groups.

|  | Control | Diabetes Patients |  |
| --- | --- | --- | --- |
| | Mean ± SD | Mean ± SD | p-value |
| **Protein** | 94.5 g/day ± 32.2 | 111.1 g/day ± 35.4 | 0.043* |
| **Carbohydrates** | 218.5 g/day ± 70.9 | 236.2 g/day ± 75.0 | 0.839 |
| **Sugar <sup>t</sup>** | 92.5 g/day ± 31.5 | 84.0 g/day ± 32.6 | 0.352 |
| **Total fat** | 73.4 g/day ± 27.2 | 85.0 g/day ± 35.8 | 0.432 |
| **Monounsaturated fat** | 34.4 g/day ± 13.2 | 40.4 g/day ± 19.0 | 0.421 |
| **Polyunsaturated fat** | 12.4 g/day ± 4.7 | 15.3 g/day ± 7.5 | 0.607 |
| **Saturated fat** | 20.4 g/day ± 8.7 | 21.9 g/day ± 8.0 | 0.421 |
| **Calories** | 1946.7 Kcal/day ± 584.1 | 2199.8 Kcal/day ± 726.4 | 0.594 |

\*Statistically significant. However, after Bonferroni correction, the adjusted p\_value was 0.387.
<sup>t</sup>Variables to which Student's T-test was applied. The remaining variables were analyzed using the Mann-Whitney test.

## Discussion

This study aimed to characterize the oral microbiota in diabetic individuals from Portugal using the 16S rRNA sequencing method to shed light on the relationship between the oral microbiome and the type 2 diabetes disease.

The predominant phyla we found in the oral microbiome of the studied samples are in line with previous studies (5, 20, 21). Likewise, the taxa frequency distribution we found in our samples follows the pattern usually observed in the oral microbiome, in which few taxa recruit most sequences (22, 23). B. J. Keijser et al. (24) study on healthy adults reported that *Prevotella, Streptococcus, and Veillonella* genera were responsible for about 50% of the total salivary microbiome, which is similar to our findings. On the other hand, we found the presence of *Gluconacetobacter* genus in 9 individuals at low frequencies (0.25-0.87%), which is not present in the Human Oral Microbiome Database (HOMD)(25), nor described, as far as we know, in other studies about the oral microbiome. Species of this genus have been found in plants (26, 27), in grapes and wine spoilage (28), and neoplastic tissue in breast cancer (29). To confirm its presence as part of the oral microbiome, other studies using oral swabs and not saliva, as the latter might contain food remains, should be performed.

The average number of taxa at the species level per individual (16) that we found was lower than what has been reported in other studies, which is around 200-600 species (13, 30). This is probably because the 16s RNA fragment we have used had a relatively low capacity to discriminate within the lowest taxonomic levels (genus and species). Sequencing of other hypervariable regions of the 16s RNA gene or the use of the whole-genome shotgun-sequencing approach might be complementary and could increase such taxonomic resolution.

Regarding the comparison of the oral microbiome of diabetics and the control group, overall, the control group showed a higher amount of taxa (202 taxa) than the diabetes group (183 taxa), which is in line with previous studies (13). Although the diabetes group had a lower richness of taxa, no significant differences were found between the two groups. Some studies reported that diabetes patients have less diversity than the control group (31) while other studies reported the opposite (32). However, in R. C. Casarin et al. (32), the diabetes patients were not controlled by medication, a factor that could explain these differences.

When we focus on particular taxa abundance differences, our results showed that *Actinobacteria* was slightly more frequent in the control group, although not statistically significant. This tendency is in line with J. Long et al. (4) findings, where *Actinobacteria* was associated with a decreased risk of diabetes. We did not find differences between both groups for the most abundant classes, *Bacilli*, and *Bacterioidia*, in contrast to A. T. M. Saeb et al. (13). These authors found that both classes were more abundant in the normal glycemic group compared to type 2 diabetic individuals. On the other hand, and in line with the results of A. T. M. Saeb et al. (13), in our study, *Gammaproteobacteria* abundance was higher but not significant in the healthy individuals, whereas *Betaproteobacteria* was significantly higher in the diabetes group (p=0.03). An interesting result was that the *TG5* genus, belonging to the *Synergistia* class, the only one that showed significant differences between groups, has been related to periodontitis (19), failed implants (33) and smoking habits (34, 35). In our findings, unexpectedly, this taxon was found in 72% of the healthy samples against only 9% of diabetes individuals. We need to take into account that the frequencies of this and other genera related to periodontitis, and thus, to diabetes (36), are vestigial in our groups, being the *TG5* just slightly higher in controls.

The almost absence of significant differences in taxa abundance frequency between the two groups, the fact that *TG5* frequency was slightly higher in controls, together with the similar microbial composition profile of control and diabetes groups according to the PERMANOVA analysis, as well as the PCoA and the hierarchical clustering, could be attributed to the fact that the diabetics were medically controlled and probably subjected to some diet restriction. These could have resulted in the lack of differences in the diet between the two groups. Therefore, it would be interesting to compare the results obtained for diabetic patients with a group of non-medically controlled diabetics. Also, we may have low statistical power due to an insufficient number of samples, and thus, it would be useful to increase the sample size.

We also investigated oral hygiene and smoking habits for its influence on microbiome diversity. The oral microbiome diversity may decrease with frequent oral hygiene habits according to M. J. Pyysalo et al. (37). This study reported that those who brushed their teeth 2 times per day presented a lower diversity than those who washed them more rarely, which is not following our findings. According to our analysis, those who wash their teeth more frequently have a higher, although not significant, diversity than those who do not wash them, probably to the fact that never washing the teeth may favor that some bacteria colonize the oral cavity preventing other species to grow. In any case, it would be necessary to perform this analysis considering a larger sample size to confirm that this tendency maintains.

As for smoking, previous studies reported that smokers tend to have a more diverse microbiota, including pathogenic taxa, than non-smokers (38, 39). Our data analysis showed that heavy smokers presented a slightly higher mean diversity index than the other two categories, but it was not significant. Perhaps the increase of the sample size and depth would help to confirm this trend. Concerning the association between type 2 diabetes and the oral microbiome, to the best of our knowledge, this study is a pioneer in the assessment of the oral microbiome diversity for a Portuguese sample of type 2 diabetes patients, and thus more difficult to interpret the obtained the data. We did not find clear differences between both groups, although it was possible to identify some taxonomic dissimilarity, which could be because the diabetes individuals were medically controlled and/or to the lack of power due to an insufficient number of samples. Thus, for future work, it would be interesting to increase the sample size, shotgun or deep sequencing and to recruit unmedicated patients. Also, the increase of taxonomic resolution at the lowest level would bring more differences in terms of species abundance. Therefore, data from other technologies like shotgun could provide further details on the oral microbiome of these samples.

## Material and Methods

### Sampling and questionnaire administration

Twenty-five patients with Diabetes Mellitus type 2 (age range 51 to 75; average age 63) and twenty-five healthy individuals (age range 42 to 74; average age 60) participated in this study. Both groups were composed of 17 males and 8 females (Table S1). Type 2 diabetes mellitus patients were approached after a general medicine or diabetic foot appointment. The study was approved by Centro Hospitalar de Vila Nova de Gaia/Espinho’s Ethics Committee and conducted according to the Declaration of Helsinki. Written informed consent was acquired from all participants.

Volunteers were ineligible if they presented less than one-third of the dentition, were under 18 years of age or had been under the influence of antibiotics less than two months before. All participants were instructed not to brush their teeth after their last meal before the saliva sample collection. In addition, each volunteer was submitted to a questionnaire, through face-to-face interviews, regarding their lifestyle, including information on smoking and oral hygiene habits, food restrictions, health status, types of medications they were taking and the period time in the case of antibiotics, was also collected. Teeth brushing habits were divided into 4 categories: washing i) more than once per day, ii) once per day, iii) once to three times per week, and iv) never washing them. Smoking habits were divided into heavy smokers, moderate smokers, and non-smokers. Additionally, a validated semiquantitative food frequency questionnaire (40, 41) was used to estimate nutrient intake to characterize the their diet as a possible influencing factor between both groups.

### DNA extraction

DNA was extracted from saliva according to D. Quinque et al. (42) and quantified using a Nano Drop-2000 spectrophotometer (Thermo Fisher Scientific Inc., MA USA).

### 16S rRNA amplification, library preparation, and sequencing

To amplify the V3-V4 hypervariable regions, the 341F/805R universal primers were used (43). A two-step Polymerase Chain Reaction (PCR) method was used to first amplify the target region and then to attach a barcode to each sample before pooling them for sequencing. The amplicon PCR contained 5 μl of DNA template at a concentration of 10 ng/μl, 5 μl of Taq PCR Master Mix kit (Qiagen), 0.4 μl of each primer (100 pmol/μl) and 3.2 μl of distilled and deionized water, in a final reaction volume of 14 μl per sample. The PCR cycling conditions were 95°C for 15 min, followed by 40 cycles of denaturation at 95°C for 30 s, annealing at 55°C for 1 min, and elongation at 72°C for 30 s. The final elongation was run at 60°C for 5 min, followed by a hold at 15°C.

The size of the amplicons was checked in a 2% agarose gel and amplicons were purified using the AMPure XP kit according to the manufacturer’s instructions. A second PCR was performed using two indices (i5 and i7) with 7 bp each. The reaction contained 5 μl of 2x Kapa HiFi Hot Start, 0.5 μl of each index, 2 μl of ultrapure water and 2 μl of first PCR product DNA, in a final volume of 10 μl per sample. PCR cycling conditions were run at 95°C for 3 min, succeeded by 10 cycles of denaturation at 95°C for 30 s, annealing at 55°C for 30 s, and elongation at 72°C for 30 s. The indexed amplicons were purified using the AMPure XP kit according to the manufacturer’s instructions followed by library quantification using a Qubit™ dsDNA BR Assay Kit (Thermo Fisher Scientific). All samples were normalized to 9 nM and pooled with 5 μl of each sample.

A TapeStation 2200 (Agilent Technologies) was used for the precise sizing and library quantification of the pool, followed by a library validation through a quantitative PCR. Finally, the pool was sequenced in an Illumina MiSeq sequencing platform, using the MisSeq v2 500-cycle sequencing kit, with a 2×250bp paired-end configuration at Novogene facilities in Hong Kong.

### Sequence processing and alignment

The obtained reads were analyzed using the Quantitative Insights into Microbial Ecology pipeline (QIIME) version 2-2019.7 (44). The paired-end sequences were processed through DADA2 (45), a quality control package in QIIME2, following the workflow: filtering, denoising, dereplication, chimera identification, and merging. After quality control, the resulting output was a feature table with the quantity of each amplicon sequence variants (ASVs) in each sample. Three samples from the diabetics group were not included in further analyses due to their small number of reads. Yet, there were no differences in age and sex between the control and diabetes groups.

For the taxonomic assignment, we used the Naïve Bayes classifier trained on the Greengenes (version 13.8) (46). The ASVs annotated as mitochondria and chloroplast were removed. One sample from the control group was also discarded because it was not reliable due to an excessive number of reads compared to the other samples and that 90% of them were assigned to a taxon not described in the oral microbiome, probably reflecting a sequencing artifact.

### Statistical analyses

The taxonomic abundance and the relative frequencies of each ASV were calculated for each sample with QIIME2, for the phylum, class, genus and species level. The differential taxa abundance between the two studied groups was evaluated through the Analysis of Composition of Microbiomes (ANCOM) as well as the Mann-Whitney U test in SPSS v.25, which was also used to compare the relative frequencies of bacteria associated with the periodontitis between the control and diabetes groups. Also, to perceive if potential differences between the microbiome of both groups could be due to the diet, nutrients consumption and energy intake were compared between groups with the Mann-Whitney U test or a Student’s T-test (in normal distribution data) with a 5% level of significance. We applied the Bonferroni correction for multiple tests.

Microbiome diversity was evaluated within individuals (alpha-diversity) with the ASVs abundance and the Shannon diversity index (47), which measures both richness and evenness, and between samples (beta-diversity) through the Bray-Curtis dissimilarity (48). For the alpha diversity analysis, we performed rarefaction curves to examine which sampling depth to use, with the QIIME2 program. All the samples were rarefied to a depth of 1973 reads. Differences in alpha diversity between the control and diabetes groups, as well as between categories of teeth brushing and smoking habits, were evaluated by the nonparametric Kruskal-Wallis H test in QIIME2.

Lastly, to investigate the microbiome composition profiles of diabetics and controls, we explored the grouping patterns with: i) a principal coordinated analysis (PCoA) based on the Bray-Curtis matrix, performed through EMPeror (49), and ii) a hierarchical clustering and a heatmap performed with the pheatmap package (50) in R (v3.5.1) (R Core Team, 2017) based on taxa abundance frequencies using only the taxa present in more than 15% of the samples. Besides this, we also tested whether the microbiome composition was statistically different between both groups using the Permutational Multivariate Analysis of Variance (PERMAVONA) analysis (52) with 999 permutations based on the Bray-Curtis matrix.

## Acknowledgments

We thank all the participants of this study, the staff from the Centro Hospitalar Vila Nova de Gaia/Espinho for the assistance when gathering the samples, Susana Lopes and Maria Magalhães for their assistance in the wet lab, and to Vítor Araújo for its thoughtful advises on data analysis. This work was supported by funds from the project NORTE-01-0145-FEDER-000007, from the Norte Portugal Regional Operational Program (NORTE2020), under the PORTUGAL 2020 Partnership Agreement, through the European Regional Development Fund (ERDF). LP-P is funded by national funds from FCT – Fundação para a Ciência e a Tecnologia, I.P. and MGV is funded through the Operational Program for Competitiveness Factors (COMPETE, EU) and UID/BIA/50027/2013 from FCT.

